# Clofoctol inhibits SARS-CoV-2 replication and reduces lung pathology in mice

**DOI:** 10.1101/2021.06.30.450483

**Authors:** Sandrine Belouzard, Arnaud Machelart, Valentin Sencio, Thibaut Vausselin, Eik Hoffmann, Nathalie Deboosere, Yves Rouillé, Lowiese Desmarets, Karin Séron, Adeline Danneels, Cyril Robil, Loïc Belloy, Camille Moreau, Catherine Piveteau, Alexandre Biela, Alexandre Vandeputte, Séverine Heumel, Lucie Deruyter, Julie Dumont, Florence Leroux, Ilka Engelmann, Enagnon Kazali Alidjinou, Didier Hober, Priscille Brodin, Terence Beghyn, François Trottein, Benoît Déprez, Jean Dubuisson

## Abstract

Drug repurposing has the advantage of shortening regulatory preclinical development steps. Here, we screened a library of drug compounds, already registered in one or several geographical areas, to identify those exhibiting antiviral activity against SARS-CoV-2 with relevant potency. Of the 1,942 compounds tested, 21 exhibited a substantial antiviral activity in Vero-81 cells. Among them, clofoctol, an antibacterial drug used for the treatment of bacterial respiratory tract infections, was further investigated due to favorable safety profile and pharmacokinetic properties. Notably, the peak concentration of clofoctol that can be achieved in human lungs is more than 20 times higher than its IC_50_ measured against SARS-CoV-2 in human pulmonary cells. This compound inhibits SARS-CoV-2 at a post-entry step. Lastly, therapeutic treatment of human ACE2 receptor transgenic mice decreased viral load, reduced inflammatory gene expression and lowered pulmonary pathology. Altogether, these data strongly support clofoctol as a therapeutic candidate for the treatment of COVID-19 patients.

**Summary:** Antivirals targeting SARS-CoV-2 are sorely needed. In this study, we screened a library of drug compounds and identified clofoctol as an antiviral against SARS-CoV-2. We further demonstrated that, in vivo, this compound reduces inflammatory gene expression and lowers pulmonary pathology.

## Introduction

The coronavirus disease 2019 (COVID-19) is having a catastrophic impact on human health as well as on the global economy, and it will continue to affect our lives for years to come (Arthi and Parman, 2021). This extraordinary situation led to the rapid development of safe and effective vaccines that have now been deployed at unprecedented scale. While COVID-19 vaccines have demonstrated their essential role in the control of the pandemic, we still lack affordable efficient therapies against SARS-CoV-2. Antivirals are indeed urgently needed to treat COVID-19 patients who have not yet been vaccinated and as a therapeutic approach to treat vaccinated people poorly protected due to waning immunity. Repurposing clinically evaluated drugs can potentially offer a fast track for the rapid deployment of treatments for this kind of emerging infectious disease. However, the first attempts of targeted repurposing strategies to treat COVID-19 patients have led to disappointing results (WHO Solidarity Trial Consortium et al., 2021). As an alternative approach, large-scale screening of clinically approved drugs through a carefully designed evaluation cascade may identify additional unanticipated therapeutic options that can be positioned for accelerated clinical evaluation (Jeon et al., 2020; Riva et al., 2020; Yuan et al., 2021). Here, we developed a high-content screen (HCS) using the Apteeus drug library (TEELibrary®), a comprehensive collection of 1,942 approved drugs, to identify molecules that exhibit antiviral activity against SARS-CoV-2. Clofoctol was selected based on its antiviral potency associated with favorable pharmacokinetic properties in human. Its further validation in a small-animal model makes it a promising candidate treatment for clinical evaluation in COVID-19 patients.

## Results

### HCS screening of a library of approved drugs

The screen was performed in Vero-81 cells, an African green monkey kidney cell line highly permissive to SARS-CoV-2 infection (Matsuyama et al., 2020). The read out was based on the cytopathic effect (CPE) of the virus as measured by propidium iodide (PI) incorporation into the nuclei of dying cells and cell quantification by nuclei staining with Hoechst 33342 (Supplementary Fig. 1**a**, Supplementary Table 1). Assay parameters, including cell seeding density, multiplicity of infection (MOI) and time points, were optimized in Vero-81 cells by measuring virus-induced CPE in a 384-well format.

To assess reproducibility of the optimized assay in a HCS configuration, we initially evaluated the effect of chloroquine (CQ), previously reported to have antiviral activity against SARS-CoV-2 in Vero Cells (Wang et al., 2020). This compound is an effective inhibitor of coronavirus entry into host cells through the endocytic pathway, however because of its lack of effect on the TMPRSS2-mediated pathway, chloroquine has been shown to be ineffective to treat COVID-19 patients. Nonetheless, this enabled us to benchmark the dynamic range of the assay with a reliable positive control. Robustness was then assessed by calculating the strictly standardized mean difference (SSMD) of each plate, with a mean of 6.87 (±2.19) for all plates (Supplementary Fig. 1**b**). We then used this experimental design to screen our drug library (Fig. 1**a**) using a non-cytotoxicity concentration of 15 μM for most compounds in the presence of low viral input (MOI = 0.01), in order to capture multicycle replication with an extended end-point measurement at 72h post-infection (Riva et al., 2020). Indeed, at this time-point, a major cytopathic effect could be observed with a strong decrease in cell number at the low MOI used in our assay. For each compound, a Robust Z-score normalized to the median of each plate was calculated for both SARS-CoV-2-induced CPE related readouts (PI measurement and Hoechst 33342 staining).

**Figure 1:**
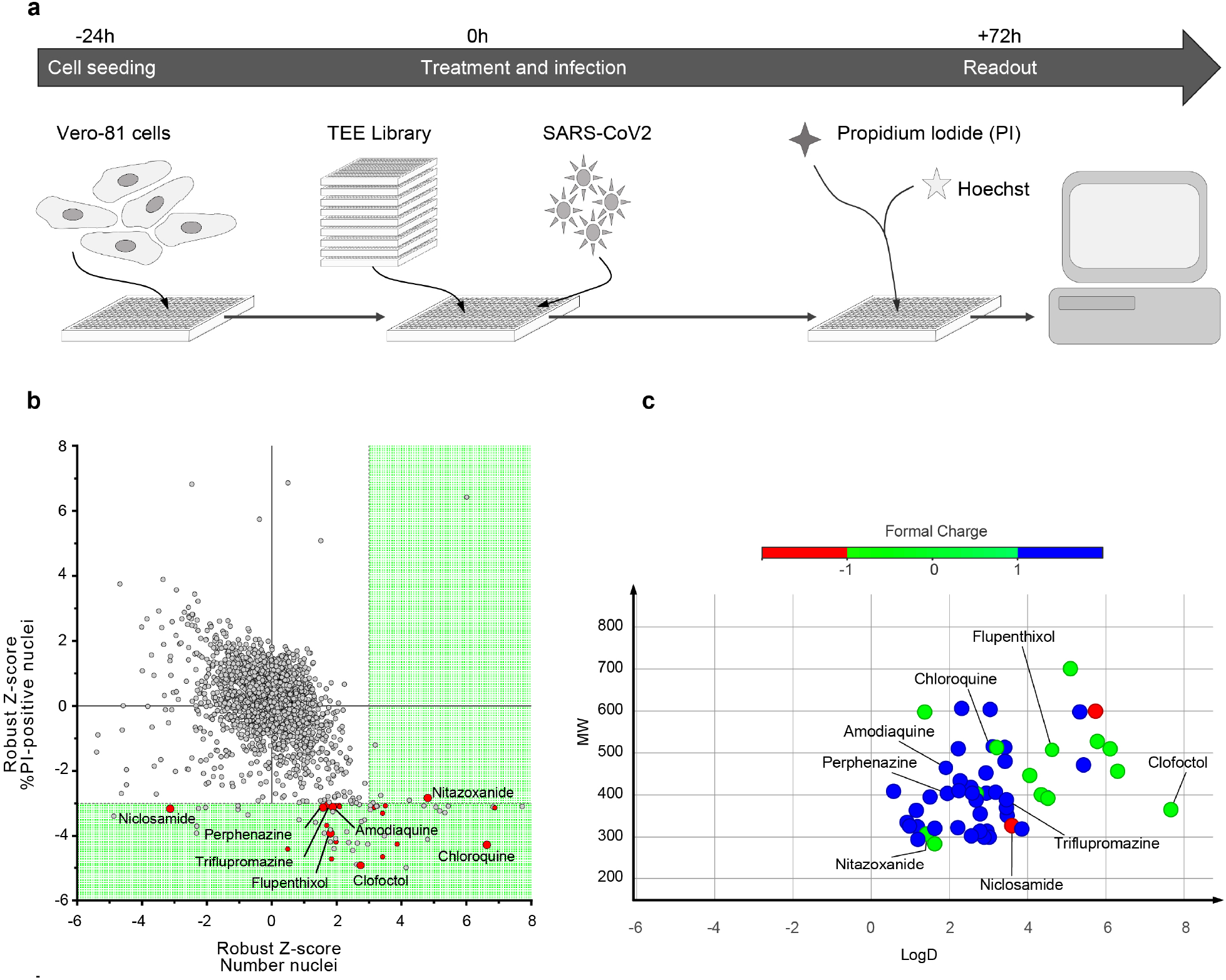
HCS screen of Apteeus TEElibrary® for the identification of anti-SARS-CoV-2 compounds. **a,** Workflow overview of the screening process. **b,** Dot-plot representations of all compounds tested based on their Robust-Z-score for both the numbers of nuclei and the percentages of PI-positive nuclei. Dotted lines are indicative of the thresholds chosen for hit selectivity (within the green area). **c**, Dot-plot representation according to the molecular weight and the LogD of the compounds of of interests. Dots are color-coded based on the ionization state at physiological pH.

Compounds exhibiting the highest levels of CPE inhibition were initially selected. Of the 1,942 tested compounds, 57 were identified to significantly decrease PI incorporation (robust Z-core < −3) or to increase the number of cells as measured with Hoechst 33342 staining (robust Z-score > 3) (Supplementary Table 2). Among these compounds, CQ, nitazoxanide, amodiaquine, triflupromazine and niclosamide were previously identified in other studies (Jeon et al., 2020; Wang et al., 2020; Lu et al., 2021; Weston et al., 2020) (Fig. 1**b**). It is worth noting that the majority of the selected compounds are basic molecules like CQ, amodiaquine, fluphenazine, trifluoperazine and triflupromazine (Fig. 1**c**). Such compounds are known to affect the endosomal internalization pathway by accumulating inside the endosomes, modifying their pH and therefore inhibiting SARS-CoV-2 by blocking its endosomal entry.

We assessed the activity of the 57 identified hits in dose-response curve (DRC) analyses using the same setting as in the screen (Supplementary Table 2). Of these, 21 showed dose-related inhibition of the PI incorporation upon infection (N=2).

Coronaviruses, including SARS-CoV-2, can use two different routes to enter their target cells. They either enter cells by endocytosis and release their genome into the cytosol after fusion of their envelope with an endosomal membrane, or they fuse their envelope directly with the plasma membrane. This latter entry route is triggered by the cell surface protease TMPRSS2 which is not expressed in Vero-81 cells (Hoffmann et al., 2020). Previously identified anti-SARS-CoV-2 compounds, like CQ and hydroxy-CQ, were shown to only block the endocytic entry route (Hoffmann et al., 2020). An additional validation on Vero-81-TMPRSS2 cells was therefore performed to discard compounds that only block the endocytic route of SARS-CoV-2 entry.

Of the 21 molecules validated twice in Vero-81 cells, the most interesting ones were retained based on a preliminary evaluation of their risk/benefit ratio in the clinic. This evaluation included a comparison of the *in vitro* potency to plasma exposure at the approved dose. Therefore, only 8 out of the 21 compounds were tested in Vero-81-TMPRSS2 cells (Supplementary Table 2). Only 3 of them exhibited a dose-dependent antiviral activity against SARS-CoV-2 in the presence or absence of TMPRSS2, indicating an antiviral effect irrespective of the entry route. These three compounds are perphenazine, nitazoxanide, and a third one, called clofoctol, that was not reported by others. More importantly, clofoctol is well distributed in tissues, particularly in lungs where its concentration is twice higher than in plasma (Del Tacca et al., 1987). Altogether, these observations prompted us to further characterize the anti-SARS-CoV-2 properties of clofoctol.

### *In vitro* validation of the antiviral activity of clofoctol

To further confirm the antiviral activity of clofoctol, SARS-CoV-2 genomic replication was measured by quantitative RT-PCR. In this assay, clofoctol exhibited an IC_50_ of 12.41 μM in Vero-81 cells and 13.51 μM in Vero-81-TMPRSS2 (Fig. 2**a**). To validate the specificity of its antiviral activity, its potential cytotoxic effect in cell culture was determined in an MTS viability assay. As shown in Fig. 2**b**, after 24h of treatment, no cytotoxic effect was observed at concentrations below 40 μM, indicating that the decreased SARS-CoV-2 replication in the presence of clofoctol was not due to a cytotoxic effect of the compound. To further characterize the antiviral activity of clofoctol, its effect on the production of infectious progeny virions was also quantified. As shown in Fig. 2**c**, a dose-dependent decrease of infectious virus production was observed in these experimental conditions, confirming the antiviral effect of clofoctol against SARS-CoV-2 with an IC_50_ of 9.3 μM and 11.59 μM in Vero-81 and Vero-81-TMPRSS2 cells, respectively.

**Figure 2:**
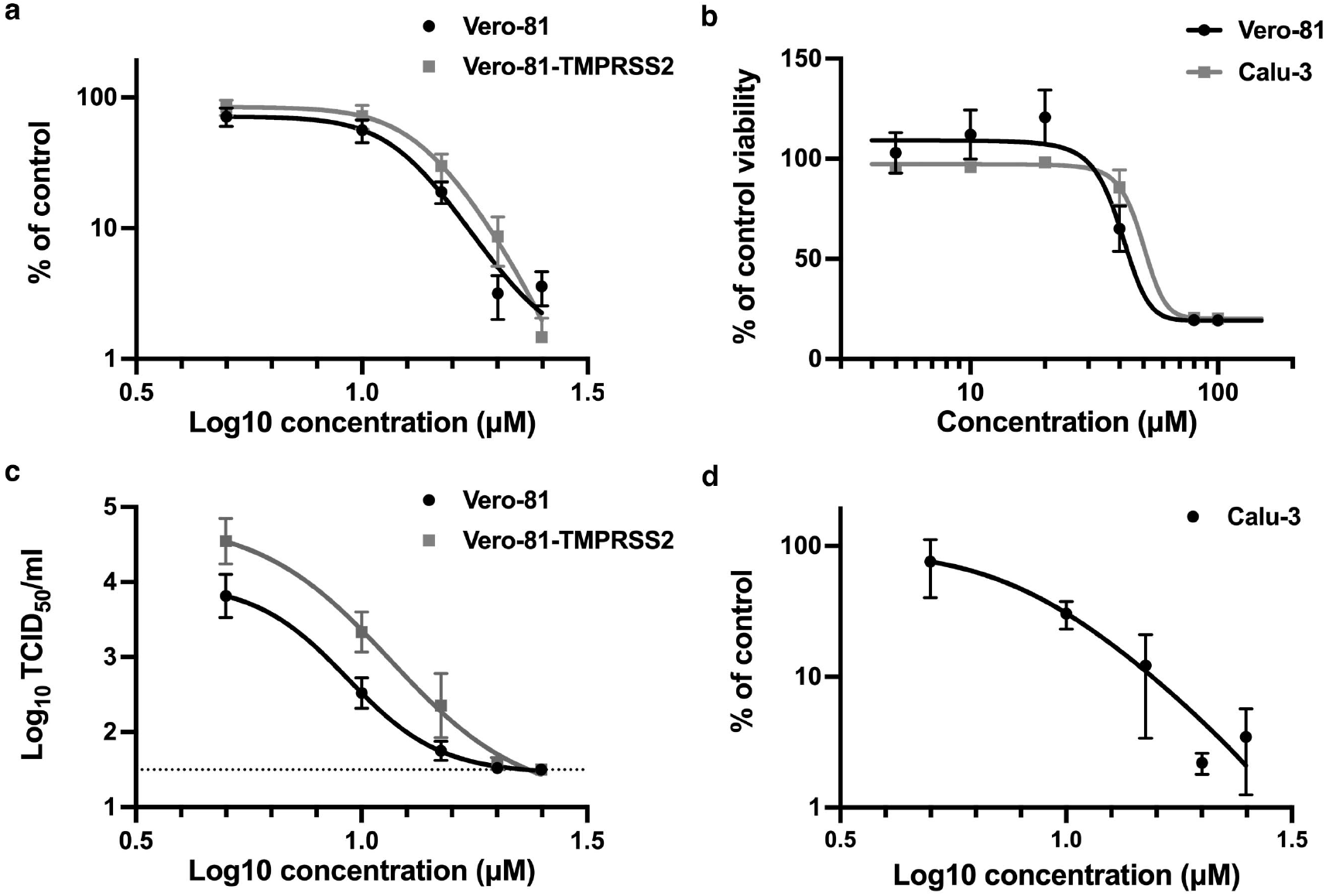
In vitro validation of the antiviral activity of clofoctol. **a**, Clofoctol inhibits the genomic replication of SARS-CoV-2. Vero-81 and Vero-81-TMPRSS2 cells were infected for 6h at an MOI of 0.25 in the presence of increasing concentrations of clofoctol. Then, total RNA was extracted and viral RNA was quantified by RT-qPCR and normalized by the amount of total RNA. Results are presented as the percentage of the viral load of the control and represent the average of seven independent experiments performed in duplicates. Error bars represent the standard error of the mean (SEM). **b**, Clofoctol is not cytotoxic in cell culture at concentrations below 40 μM. Vero-81 cells and Calu-3 cells were cultured in the presence of given concentrations of clofoctol. Cell viability was monitored using the MTS-based viability assay after 24 hours of incubation. **c**, Clofoctol inhibits the production of progeny virions. Vero-81 and Vero-81-TMPRSS2 cells were infected with SARS-CoV-2 at a MOI of 0.25. After 1h, the inoculum was removed and the cells were washed with PBS prior treatment with clofoctol. Cells were then further incubated for 16h. Thereafter, supernatants were collected and the amounts of secreted infectious virus were quantified. The dotted-line represents the limit of detection (1.5 TCID_50_/mL). These data represent the average of three independent experiments (N=3). Experiments were performed in duplicate for each condition. **d**, Clofoctol inhibits SARS-CoV-2 replication in Calu-3 cells. Calu-3 cells were infected at a MOI of 0.25 in the presence of increasing concentrations of clofoctol for 24h. Then, total cellular RNA was extracted and viral RNA was quantified by RT-qPCR. Results are presented as the percentage of the viral load of the control and represent the average of three independent experiments performed in duplicates. Error bars represent the standard error of the mean (SEM)

Vero cells are the cells of choice to efficiently grow SARS-CoV-2 in culture and therefore to screen large libraries of compounds for rapid identification of antivirals. However, as these cells are from monkey origin, the human cell line Calu-3, derived from a lung adenocarcinoma, previously shown to be permissive to SARS-CoV-2 (Hoffmann et al., 2020) was also used to validate our observations. As shown in Fig. 2**d**, a dose-dependent decrease of viral RNA production was also observed in infected Calu-3 cells treated with clofoctol, at concentrations that did not exhibit a cytotoxic effect (Fig. 2**b**). In this cell line, clofoctol exhibited an IC_50_ of 7.9 μM.

### Clofoctol inhibits the translation of SARS-CoV-2 viral RNAs

The life cycle of a virus can be divided into three major steps: (1) entry, (2) translation/replication and (3) assembly/release. To determine at which step clofoctol inhibits SARS-CoV-2, the compound was added either before infection, during virus entry, post-inoculation or throughout all the steps. Remdesivir, an inhibitor of the viral polymerase (Lo et al., 2020), and CQ were used as control antivirals affecting viral replication or entry, respectively. As shown in Fig. 3**a**, remdesivir inhibited infection only when added after the entry step, whereas CQ was only efficient when added at the entry step. Clofoctol inhibited SARS-CoV-2 mainly at the post-inoculation step, although it had also a mild effect at the entry step. These data suggest that the translation/replication step is likely the major target of clofoctol. To further characterize the post-entry inhibitory effect of clofoctol, a time-of-addition experiment was also performed in parallel with clofoctol, remdesivir and CQ. In this experiment, the compounds were added before infection or at different times post-infection. As shown in Fig. 3**b**, CQ was only efficient when added prior to infection or during the inoculation step, whereas remdesivir and clofoctol remained effective when added up to three or four hours post-inoculation, respectively. To further exclude a potential effect of clofoctol on SARS-CoV-2 entry, we used retroviral particles pseudotyped with the SARS-CoV-2 Spike (S) glycoprotein (SARS2pp). These are retroviral cores carrying SARS-CoV-2 S glycoprotein in their envelope and a minigenome containing a luciferase reporter gene. In this context, only the early steps of the viral life cycle (i.e., virus interaction with receptors, uptake, and fusion) are SARS-CoV-2 specific, whereas all later steps are dependent on retroviral nucleocapsid elements. Clofoctol did not show any inhibitory effect on SARS2pp entry (Fig. 3**c**), indicating that it does not inhibit the cellular entry of SARS-CoV-2. Of note, clofoctol was also active against another coronavirus which is mildly pathogenic, HCoV-229E, and more importantly against the D614G, the B.1.1.7 and the B.1.351 variants of SARS-CoV-2 (Supplementary Fig. 2), indicating that its antiviral activity is conserved across different clades of SARS-CoV-2 and coronavirus species.

**Figure 3:**
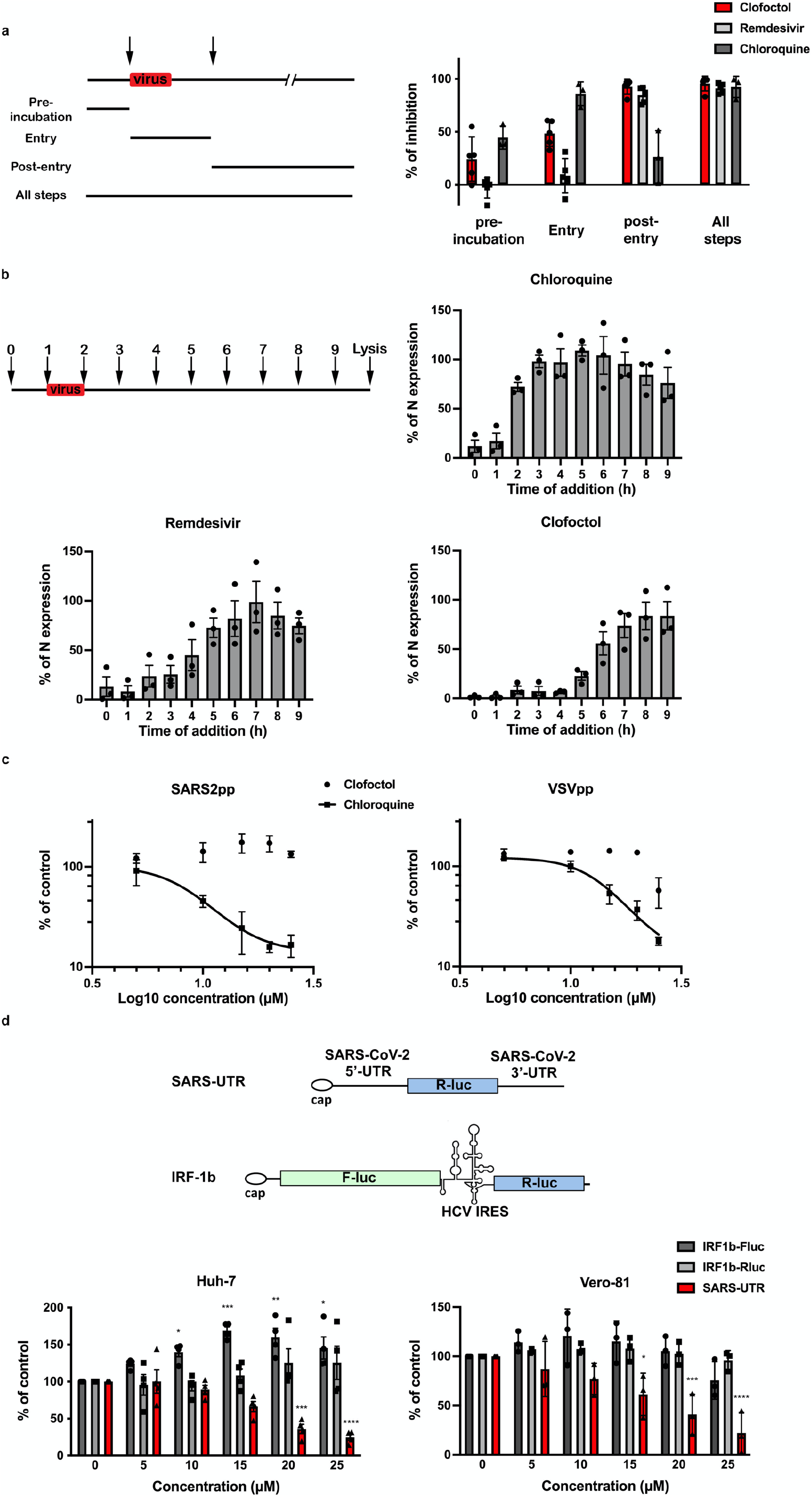
Clofoctol targets the translation step of SARS-CoV-2 life cycle. Clofoctol targets the translation step of SARS-CoV-2 life cycle. **a**, Clofoctol is mainly efficient at a post-entry step. Vero-81 cells were infected with SARS-CoV-2 at an MOI of 0.25. Clofoctol, remdesivir or CQ were added at a concentration of 15 μM either before infection, during virus entry, post-inoculation or throughout the steps as indicated. At 16h post-infection, cells were fixed with 4% paraformaldehyde and processed to detect the proportion of infected cells. Therefore, they were immunostained to allow for the detection of the viral double-stranded RNA and nuclei were detected by Hoechst staining to count the total number of cells. Results are presented as the percentage of infection inhibition and represent the average of five independent experiments. **b**, Time-of-addition experiment. Vero-81 cells were infected at an MOI of 0.5 for 1h. 15 μM of clofoctol, remdesivir or CQ were added every hour starting 1h before inoculation. Cells were lysed 8h after the end of the inoculation in Laemmli loading buffer and the amount of N protein was detected in immunoblot. Results are presented as the percentage of N protein expression relative to that in non-treated cells (CTL) and represent the average of three independent experiments. Error bars represent the standard error of the mean (SEM). **c**, Clofoctol does not inhibit SARS-CoV-2 entry. Huh-7 cells expressing ACE2 receptor were infected with SARS2pp or pseudoparticles containing the envelope glycoprotein of the vesicular stomatitis virus (VSV) used as a control (VSVpp) for 3 hours in the presence of increasing concentrations of clofoctol or CQ. At 48 hours post-infection, cells were lysed to quantify luciferase activity. The results are expressed in % of the controls of three independent experiments. The experiments were performed in triplicate (n=3) (in each condition). **d**, Clofoctol inhibits viral RNA translation. Schematic representation of the reporter construct expressing the *Renilla* luciferase placed between the 5’-UTR and the 3’-UTR of the SARS-CoV-2 genomic RNA and the control bicistronic construct containing the firefly luciferase sequence under the control of a cap structure, followed by the *Renilla* luciferase under the control of hepatitis C virus (HCV) IRES. Huh-7 or Vero-81 cells were electroporated with *in vitro* transcribed RNA. Cells were lysed after 8h and luciferase activities were recorded. The results are expressed in % of the controls of three independent experiments. The experiments were performed in quadruplicate (n=4)(in each condition). Two-way ANOVA followed by the Dunnett’s multiple comparisons test was performed for statistical analysis (*p < 0.05; **p < 0.01; ***p < 0.001).

For positive-stranded RNA viruses like coronaviruses, translation of the viral genome is the step that immediately follows virus entry. To determine whether this step is affected by clofoctol treatment, we used a reporter construct expressing the *Renilla* luciferase introduced between the 5’-UTR and the 3’-UTR of the SARS-CoV-2 genomic RNA. As a control, we used a bicistronic construct containing the *Firefly* luciferase sequence under the control of a eukaryotic mRNA 5’ cap structure, followed by the *Renilla* luciferase sequence under the translational control of the hepatitis C virus (HCV) IRES (Goueslain et al., 2010). To avoid a potential effect of clofoctol on the transcription of the reporters from plasmid DNA, *in vitro*-transcribed capped RNAs were transfected by electroporation into Vero-81 or Huh-7 cells. After 8h of clofoctol treatment, a dose-dependent inhibition of luciferase activity was observed with the SARS-CoV-2 UTRs-based construct in Vero-81 and Huh-7 cells, but not for the control bicistronic construct (Fig. 3**d**), indicating that clofoctol specifically inhibits the translation of an mRNA containing the UTRs of the SARS-CoV-2. This suggests that clofoctol has the potential to inhibit the translation of genomic as well as sub-genomic SARS-CoV-2 RNAs.

### Clofoctol inhibits SARS-CoV-2 replication *in vivo* and lowers inflammation in lungs

To investigate the potential antiviral activity of clofoctol against SARS-CoV-2 *in vivo*, we took advantage of transgenic C57BL/6 mice expressing the human ACE2 receptor (K18-hACE2 mice) (Golden et al., 2020). Before testing the potential antiviral activity of clofoctol, pharmacokinetic experiments were performed in female C57BL/6 mice. To this end, clofoctol was injected intraperitoneally (i.p.) at 62.5 mg/kg to reach a lung concentration close to that achieved in humans at approved posology. Mice were sacrificed at 30 min, 1h, 2h and 4h after i.p. administration of clofoctol. As early as 30 min after injection, clofoctol reached concentrations up to 61 μM in the lungs and remained above this level for almost 4h (Fig. 4**a**, *left panel*), whereas it remained at a concentration seven times lower in the plasma. According to its expected half-life in the lungs, clofoctol concentration was anticipated to remain above its *in vitro* measured IC_50_ (IC_50_ ~10μM) for more than 7 consecutive hours. It was then decided to treat the mice twice daily to maintain a lung concentration close to 60 μM. In this setting, clofoctol concentration reached 67 μM in the lungs, 1h after the fourth administration (Fig. 4**a**, *right panel*).

**Figure 4:**
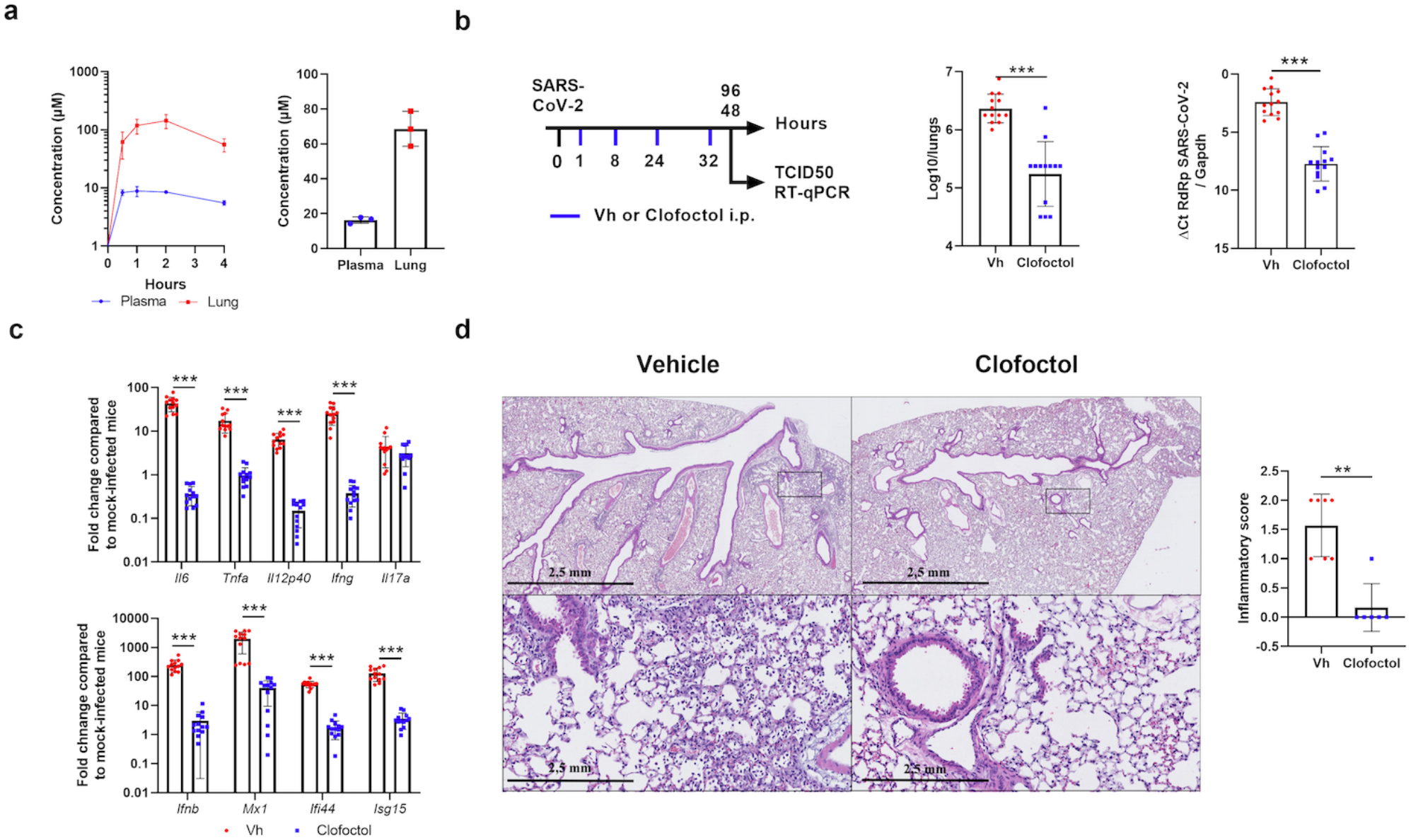
Pharmacokinetics and antiviral properties of clofoctol in a mouse model of COVID-19. **a**, Pharmacokinetics characterization of clofoctol in mice. *Left* panel, 8-10 week-old female C57BL/6J mice were treated i.p. with a single dose of clofoctol (62.5mg/kg) and were sacrificed at different time points thereafter. *Right* panel, Clofoctol was inoculated twice daily during two days and mice were sacrificed 1h after the last injection. Clofoctol concentrations in lungs (n=3/time point, 3 samples/lung) and plasma (n=3/time point, 2 technical replicates) are depicted. **b**-**d**, Effects of clofoctol treatment on SARS-CoV-2 infection in K18-hACE2 transgenic C57BL/6J mice. **b**, *Left panel*, Scheme of the experimental design in which the effects of clofoctol was assessed in mice. Mice were treated i.p. with clofoctol (62.5mg/kg) or vehicle 1h and 8h after i.n. inoculation of SARS-CoV-2 (5×10^2^ TCID_50_ per mouse) and treated again twice at day 1 post-infection. Animals were sacrificed at day 2 and day 4 post-infection. The viral load was determined by titration on Vero-E6 cells (*middle panel*) and by RT-qPCR (*right panel*) (day 2 post-infection). **c**, mRNA copy numbers of genes were quantified by RT-qPCR. Data are expressed as fold change over average gene expression in mock-treated (uninfected) animals (day 2 post-infection). **d**, Lung sections were analyzed at day 4 post-infection. Shown are representative lungs (hematoxylin and eosin staining). *Lower panels*, enlarged views of the area circled in black in *upper* panels. Blinded sections were scored for levels of pathological severity. The inflammatory score is depicted. **b**-**c**, Results are expressed as the mean ± SD (n=13 for panels **b** and **c** and n=6-7 for panel **d**). Significant differences were determined using the Mann-Whitney U test (**p < 0.01; ***p < 0.001).

Because of this favorable pharmacokinetic profile in mice, we decided to test clofoctol in K18-hACE2 transgenic mice. Female mice were inoculated intranasally (i.n.) with a lethal dose (5×10^2^ TCID_50_) of a clinical SARS-CoV-2 isolate. The animals were then injected intraperitoneally with clofoctol at 1h and 8h post-infection. This treatment was repeated the day after infection and some of the mice were sacrificed at day 2 post-inoculation (Fig. 4**b**, *left panel*). As compared to untreated animals, the infectious viral load detected in the lungs of clofoctol-treated mice was reduced by more than 1.1 log_10_ at day 2 post-infection (Fig. 4**b**, *middle panel*). Analysis of viral RNA yields by RT-qPCR confirmed the reduced viral load in clofoctol-treated animals (Fig. 4**b**, *right panel*). Male K18-hACE2 transgenic mice are more susceptible than females to SARS-CoV-2 infection (Golden et al., 2020). Clofoctol treatment similarly reduced the viral load in the lungs of male K18-hACE2 transgenic mice (Supplementary Fig. 3**a**).

We then investigated whether the decrease in viral load would have positive effects on lung inflammation. Remarkably, the expression of transcripts encoding IL-6, TNFα, IL12p40, IFNβ, IFNγ and the interferon-stimulated genes (ISG) Mx1, Ifi44 and ISG15 was markedly reduced in clofoctol-treated mice, in contrast with that of IL-17A (Fig. 4**c**). Similar results were also observed in male K18-hACE2 transgenic mice (Supplementary Fig. 3**b**). At day 4 post-inoculation, relative to controls, mice treated during the first 2 days with clofoctol still showed a lower viral load and a decreased expression of transcripts encoding inflammatory markers (supplementary Fig. 4**a**, Fig. 4**b** and data not shown). At this time point, SARS-CoV-2 infection was associated with reduced expression of genes encoding markers of epithelial barrier function, including the tight-junction protein Zonula Occludens-1 (ZO-1) and occludin. Interestingly, the drop of these transcript levels induced by SARS-CoV-2 infection was significantly reduced in clofoctol-treated animals (supplementary Fig. 4**c**).

Lastly, we assessed the impact of clofoctol treatment on lung pathology at day 4 post-infection. In vehicle-treated animals, a mild multifocal broncho-interstitial pneumonia was observed (Fig. 4**d**, *upper left panel*). Signs of moderate inflammation, with the presence of neutrophils, macrophages and a few lymphocytes, were observed within alveolar lumens, inter-alveolar septa, and perivascular spaces which was accompanied by minimal perivascular edema (Fig. 4**d**, *lower left panel*). Slight vascular congestion and discrete intra-alveolar hemorrhages were also detected (not shown). In stark contrast, only a minimal interstitial inflammation was observed in clofoctol-treated mice (Fig. 4**d**, *upper right panel*), with a limited presence of macrophages and lymphocytes within inter-alveolar septa and little vascular congestion (Fig. 4**d**, *lower right panel*). We conclude that, at doses that produce lung concentrations close to those observed in human patients treated at the approved dose, clofoctol treatment in mice just after infection lowers SARS-CoV-2 replication and reduces lung pathological features associated with this viral infection.

## Discussion

In this study, we report the high-throughput screening of ~2,000 drugs, approved for human use, for their potential activity against SARS-CoV-2. Our data identify clofoctol as a promising antiviral candidate for the treatment of COVID-19 patients. This antibacterial drug was developed in the late 1970s. Its efficacy has been demonstrated for the treatment of *Streptococcus pneumoniae* - the leading cause of bacterial pneumonia worldwide - and *Staphyloccus aureus* (Danesi and Del Tacca, 1985; Ghilardi and Casani, 1985). The drug was marketed in France until 2005 under the trade name Octofène® and is still prescribed in Italy under the trademark GramPlus®. Mechanistically, clofoctol inhibits bacterial cell wall synthesis and induces membrane permeabilization (Yablonsky, 1983; Yablonsky and Simonnet, 1982). Along with its bactericidal activity, clofoctol was recently shown to also inhibit protein translation and to impair tumor cell growth (Hu et al., 2019; Wang et al., 2014a). As such, clofoctol could be useful to treat some cancers and possibly other diseases (Bailly and Vergoten, 2021).

Among the 2,000 drugs tested, clofoctol emerged as the most promising compound to inhibit SARS-CoV-2 replication in our experimental settings. Our data show that it can contribute to inhibition of SARS-CoV-2 propagation by blocking translation of viral RNA. However, we cannot exclude other effects of clofoctol on SARS-CoV-2 replication. The inhibition of translation by clofoctol could be due to the activation of the unfolded protein response (UPR) pathways. Clofoctol has indeed been reported to induce endoplasmic reticulum (ER) stress and to activate all three UPR pathways, i.e. the inositol requiring enzyme 1 (IRE1), the double stranded RNA-activated PK-like ER kinase (PERK), and the activating transcription factor 6 (ATF6) (Wang et al., 2014b). Although UPR activation is observed during SARS-CoV-2 infection (Echavarría-Consuegra et al., 2021), chemical activation of UPR by thapsigargin has been shown to inhibit coronavirus replication, including SARS-CoV-2 (Shaban et al., 2021). Furthermore, modulating the PERK-eIF2α pathway can inhibit the replication of the transmissible gastroenteritis porcine coronavirus (Xue et al., 2018). Similarly, triggering the UPR with 2-deoxy-D-glucose inhibits the replication of another coronavirus, the porcine epidemic diarrhea virus (Wang et al., 2014b). Whether the clofoctol-induced inhibition of SARS-CoV-2 translation is linked to UPR activation will be the focus of further investigation.

Previous pharmacokinetic studies indicate that clofoctol is well absorbed by rectal administration, and can rapidly expose lung tissues (Del Tacca et al., 1987; Alessandrì et al., 1986). Of interest, as early as 90 minutes after rectal administration, the peak concentration of clofoctol that can be achieved in human lungs is more than 20 times higher than its IC_50_ measured in Vero-81 cells. In our experimental conditions, clofoctol was also detected in mice lungs at a peak concentration reaching approximately tenfold its IC_50_. Notably, upon two days of treatment with doses allometrically similar to those approved for human treatment, its concentration in the lungs remained far above the IC_50_ measured *in vitro*. Importantly, we demonstrate here that clofoctol treatment decreased the viral load in the lungs and drastically reduced pulmonary inflammation. These *in vivo* data, as well as the rapid onset of action expected in human pharmacokinetics, strongly support clofoctol as a therapeutic candidate for the treatment of COVID-19 patients. An ongoing phase 2/3 placebo controlled clinical trial should further validate the therapeutic potential of this compound in the early phase of COVID-19.

Together with its antiviral effects, clofoctol abrogated lung inflammation. To the best of our knowledge, the anti-inflammatory effect of clofoctol has never been reported before. Together with its effect on UPR pathways, clofoctol is known to interact with different targets including (i) the Cdc7/Dbf4 protein kinase complex, which regulates the initiation of DNA replication and (ii) the upstream-of-N-Ras protein (UNR), a highly conserved RNA-binding protein known to regulate gene expression. Of interest, by binding to UNR, clofoctol activates the transcription factor Kruppel-like factor 13 (KLF13)(Wang et al., 2014a), known as a tumor suppressor gene and as a regulator of T cell differentiation (Fernandez-Zapico et al., 2011; Jiang et al., 2015). Whether the UNR/KLF13 pathway triggered by clofoctol plays a role in decreasing inflammation during SARS-CoV-2 infection deserves further investigation. Additional functional studies are urgently needed to assess the global effect of clofoctol on COVID-19 pathology.

In conclusion, the antiviral and anti-inflammatory properties of clofoctol, associated with its safety profile and unique pharmacokinetics make a strong case for proposing clofoctol as an affordable therapeutic candidate for the treatment of COVID-19 patients.

## Methods

### Data reporting

No statistical methods were used to predetermine sample size. Compounds were spotted in a randomized order on the plates during the primary screen. All the other experiments were not randomized. Investigators were blinded to allocation during the primary screen and the corresponding validation, during both assay performance and outcome assessment. For all the other assays, the investigators were not blinded.

### Cells and viruses

Vero-81 cells (ATCC, CCL-81), Vero-E6 cells (ATCC, CRL-1586), Huh-7 cells(Nakabayashi et al., 1982) and HEK293T/17 cells (ATCC, CRL-11268) were grown at 37°C with 5% CO_2_ in Dulbecco’s modified eagle medium (DMEM, Gibco) supplemented with 10 % heat-inactivated fetal bovine serum (FBS, Eurobio). Calu-3 cells (Clinisciences, EP-CL-0054) were grown in minimum essential medium (Gibco, MEM) supplemented with glutamax (Gibco) and 10% heat-inactivated FBS.

Lentiviral vectors expressing TMPRSS2 were produced by transfection of HEK293T cells with pTRIP-TMPRSS2, phCMV-VSVG and HIV gag-pol in the presence of Turbofect (Life Technologies) according to the manufacturer’s instruction. Supernatants were collected at 48h post-transfection and used to transduce Vero-E6 cells.

The BetaCoV/France/IDF0372/2020 strain of SARS-CoV-2 was supplied by the French National Reference Center for Respiratory Viruses hosted by Institut Pasteur (Paris, France). The hCoV-19_IPL_France strain of SARS-CoV-2 (NCBI MW575140) was also used for *in vivo* experiments. All SARS-CoV-2 viruses, including the variants B.1.1.7 and B.1.351 were propagated in Vero-E6 cells expressing TMPRSS2 by inoculation at MOI 0.01. Cell supernatant medium was harvested at 72h post-infection and stored frozen at −80 °C in small aliquots. All experiments were conducted in a biosafety level 3 (BSL3) laboratory.

### Chemical libraries

The TEELibrary® was built and supplied by APTEEUS company. It was in its version n°4 and counted 1,942 small organic molecules approved for a use in human and selected within national and international drug repositories. It is mainly composed of active pharmaceutical ingredients (>90%) and it covers 85% of the Prestwick FDA approved collection. All molecules have been dissolved in an appropriate bio-compatible solvent (DMSO or water with adjusted pH), at a concentration compatible with the testing on living cells. The majority of them are prepared at a 10mM concentration in DMSO. CQ diphosphate was purchased from Sigma-Aldrich (Dorset, England and St. Louis, MO). CQ diphosphate was diluted to a final concentration of 10 mM in water. Clofoctol was purchased from Sigma-Aldrich (C2290) or provided by Chiesi company.

### Drug screening assay

One day prior to infection, Vero-81 cells were seeded in black 384-well μClear® plates (Greiner Bio-One), at a density of 3,000 cells per well in 30 μl DMEM, supplemented with 10% FBS and 1X Penicillin-Streptomycin solution (Gibco), using MultiDrop Combi® Reagent dispenser (ThermoFischer Scientific). The next day, compounds from the TEELibrary® were first dispensed into the 384-well plates, using an Echo 550 Liquid Handler (Labcyte). To identify the compounds of interest, they were tested at a final compound concentration that usually does not induce cytotoxicity, most of them at 15 μM. On each plate, five 3-fold serial dilutions of CQ diphosphate ranging from 0.15 μM to 15 μM were added in six replicates, as a control compound of viral inhibition (positive controls). Eleven control virus wells devoid of compound and scattered over the plate, were supplemented with 0.15% DMSO or 0.15% H_2_O (negative controls), respectively. Cells were infected by adding 10 μL of SARS-CoV-2 per well at a MOI of 0.01 in 10% FBS-containing medium, using a Viafill Rapid Reagent Dispenser (Integra). The plates were then incubated at 37° with 5% CO_2_. At 3 days post-infection, cells were stained with 10 μg/mL Hoechst 33342 dye (Sigma-Aldrich) and 1 μg/mL PI (ThermoFischer Scientific) for 30 min at 37°C for CPE quantification by high-content imaging.

### Dose response curves and hit validations

The selected hits were further validated in a 6-point dose-response confirmation assay. One day prior to infection, Vero-81 cells were seeded in 384-well plates, as previously described. The next day, six 3-fold serial dilutions of compounds (0.15 to 45 μM, in duplicate) were first added to the cells. Ten μL of virus diluted in medium was then added to the wells. On each plate, twenty-six virus control wells distributed over the plates were supplemented with 0.15% DMSO and H_2_O, respectively. CQ diphosphate was added as a control compound, at six 3-fold serial dilutions (0.15 μM to 45 μM, in duplicate). Plates were incubated for 3 days at 37°C prior to staining and CPE quantification by high-content imaging.

### Image acquisition

Image acquisitions were performed on a high-resolution automated confocal microscope (Opera, PerkinElmer) using a 10x air objective for cellular infection assay. Hoechst 33342-stained nuclei were detected using the 405 nm excitation laser (Ex) with a 450/50-nm emission filter (Em). Red signal, corresponding to PI-stained nuclei from dead cells, was detected using Ex at 561 nm and Em at 600 nm. A set of 3 fields was collected from each well.

### Image-based analysis

For total cell and dead cell detection, images from the automated confocal microscope were analyzed using multi-parameter scripts developed using Columbus system (version 2.3.1; PerkinElmer) (Supplementary Table 1). A segmentation algorithm was applied to detect nuclei labeled by Hoechst 33342 (blue) and determine total nuclei number. Briefly, a mask was first determined from input image, using the intensity threshold of Hoechst dye signal to create a region of interest corresponding to Hoechst-stained population. The nuclei segmentation was then performed using the algorithm “Find Nuclei”, as described previously(Song et al., 2017). Morphology properties, as area and roundness, could be used to exclude smaller objects not corresponding to nuclei. The total number of cells was quantified as Hoechst-positive nuclei. Red fluorescence signal intensities in the previous selected nuclei were quantified and used for the selection of PI positive (PI+) and negative (PI−) nuclei. Subsequently, population of dead (PI+) and viable (PI-) cells were determined. The percentage of PI+ cells was calculated for each compound to select drugs having an effect on the decrease of cell death, corresponding to infection or viral replication inhibition.

### Dose-response validation in different cell lines

Vero-81, Vero-81-TMPRSS2 or Calu-3 cells were infected in duplicates at a MOI of 0.25 in the presence of increasing concentrations of clofoctol, ranging from 0 to 25 μM, and incubated either for 6h (Vero-81 cells) or 24h (Calu-3 cells). Then total RNA was extracted by using the Nucleospin RNA kit (Macherey Nagel) as recommended by the manufacturer. Genome quantification was performed as described (Eymieux et al., 2021).

### Viral secretion

Vero-81 and Vero-81-TMPRSS2 cells were infected at a MOI of 0.25 for 1h, then the cells were rinsed 3 times with PBS and further incubated in the presence of increasing concentrations of clofoctol for 16h. Each condition was performed in duplicates. Cell supernatants were collected and viral titer were measured by the TCID_50_ method.

### Pseudoparticles infection

Retroviral Murine leukemia virus particle were pseudotyped with the SARS-CoV-2 Spike (BetaCoV/France/IDF0372/2020 strain) or the glycoprotein of the vesicular stomatitis virus (VSV-G). Briefly, HEK293T cells were co-transfected with a plasmid encoding Gag-Pol (pTG-Gag-Pol), a plasmid encoding the envelope glycoprotein and a plasmid containing a minigenome with a *Firefly* luciferase reporter gene. After 48h of incubation, cell supernatants were collected, filtered and used to transduce Huh-7 cells expressing human ACE2 in the presence of increasing concentrations of clofoctol or CQ. Transduced cells were lysed 48h later and luciferase activity was measured by using the luciferase assay system (Promega).

### Time-of-addition experiment

Vero-81 cells were plated in 24-well plates and infected for 1h at a MOI of 0.5. Clofoctol, remdesivir or CQ were added to the cells at a concentration of 15 μM every hour starting one hour before inoculation. At 8h post-infection, the cells were lysed in non-reducing Laemmli loading buffer. Proteins were separated onto a 10% SDS-polyacrylamide gel electrophoresis and transferred on nitrocellulose membranes (Amersham). Membrane-bound N proteins were detected with a rabbit polyclonal antibody (Novus) and a horseradish peroxidase-conjugated secondary antibody (Jackson Immunoresearch). Detection was carried out by chemoluminescence (Pierce) and signals were quantified by using the gel quantification function of ImageJ. The experiment was repeated 3 times in duplicates.

### Immunofluorescence

Vero-81 cells were plated onto glass coverslips. The day after, the cells were infected for 1h with SARS-CoV-2 at a MOI of 0.25. Clofoctol, remdesivir or CQ were added at 15 μM at different steps of the infection. The cells were either incubated 1h before inoculation (pre-incubation) or during the inoculation and for 1h after virus removal (entry step) or starting 1h after the inoculation until cell fixation (post-entry). Additional conditions with the compounds present during the whole experiment were also included as well as controls with DMSO or H_2_0. Cells were incubated for 16h after infection and fixed with 4% paraformaldehyde. Then, cells were permeabilized for 5 min with 0.1% Triton X-100 in PBS and blocked for 30 min with 5% goat serum in PBS. Infected cells were detected by using an anti-dsRNA (J2 monoclonal antibody, Scicons) diluted in blocking buffer to detect the presence of replicating SARS-CoV-2 virus as previously determined (Eymieux et al., 2021). After a 30-min incubation, cells were rinsed 3 times for 5 min in PBS and incubated for 30 min with a cyanine 3-conjugated goat anti-mouse secondary antibody (Jackson Immunoresearch) and DAPI (4′,6-diamidino-2-phenylindole).

The coverslips were rinsed with PBS 3 times for 5 min followed by a final water wash before mounting on microscope slides in Mowiol 4-88 containing medium. Images acquisitions were performed with an EVOS M5000 imaging system (Thermo Fischer Scientific) equipped with a 10X objective and light cubes for DAPI and RFP. The total number of cells was determined by counting the number of nuclei and the number of infected cells was determined by counting dsRNA-positive cells. The experiment was performed three times.

### Viability assay

Vero cells, Huh-7 cells or Calu-3 cells were plated in 96-well plates and were then incubated the next day in 100 μl of culture medium containing increasing concentrations of clofoctol for 24h. An MTS [3-(4,5-dimethylthiazol-2-yl)-5-(3-carboxymethoxyphenyl)-2-(4-sulfophenyl)-2H-tetrazolium]-based viability assay (CellTiter 96 aqueous nonradioactive cell proliferation assay, Promega) was performed as recommended by the manufacturer. The absorbance of formazan at 490 nm is detected using an enzyme-linked immunosorbent assay (ELISA) plate reader (ELx808, BioTek Instruments, Inc.). Each measure was performed in triplicate.

### Analysis of the effect of the drug on translation

A plasmid containing a synthetic gene encompassing the 5’-UTR (nucleotides 1-265) and the 3’UTR (nucleotides 29675-29903) of SARS-CoV-2 isolate Wuhan-Hu-1 (Genebank NC_045512.2) separated by two head-to-tail BbsI sites was produced by GeneCust. The coding sequence of *Renilla* luciferase amplified by PCR using primers containing BbsI sites was inserted between both UTRs by ligation of BbsI-restricted PCR and plasmid. In this way, the coding sequence of the luciferase was inserted between the UTRs without leaving an extra nucleotide in between. The plasmid was linearized by NsiI restriction, and the linearized DNA was then used as a template for *in vitro* transcription with the mMESSAGE mMACHINE T7 kit from Thermofischer Scientific, as recommended by the manufacturer. *In vitro*-transcribed capped RNA was delivered to Vero-81 and Huh-7 cells by electroporation. Cells were cultured for 8h in the presence of increasing concentrations of clofoctol. *Renilla* luciferase activities were measured with a *Renilla* luciferase assay from Promega. As a control, we used a bicistronic construct containing the *Firefly* luciferase sequence under the control of a cap structure, followed by the *Renilla* luciferase under the control of hepatitis C virus (HCV) IRES. *Firefly* and *Renilla* luciferase activities were measured with a dual-luciferase reporter assay system from Merck Millipore as previously reported (Goueslain et al., 2010).

### Pharmacokinetic study

Clofoctol diluted in 1.75% final Kolliphor® RH40 (07076, Sigma) and 1.4% final ethanol in a sodium chloride solution (0.9%) was used for intraperitoneal (i.p.) injection (62.5 mg/kg in females and 50 mg/kg in males). The concentration of clofoctol in plasma and lungs was measured at different time points post-clofoctol injection. Plasma samples and lung tissues were collected and treated with absolute ethanol, in a ratio of 1 to 10 and 1 to 50, respectively. Lung tissues were homogenized with a mechanical lysis system (Tissue Lyzer II). Supernatants were obtained by centrifugation before injection in LC-MS/MS. Samples were analysed using UPLC system Acquity I Class (Waters), combined with a triple quadrupole mass spectrometer Xevo TQD (Waters). The column, placed at 40°C, was an Acquity BEH C8 50*2.1mm, 1.7μm column (Waters) and the following mobile phases were used: 5mM ammonium formate pH 3.75 in water, as solvent (A) and 5 mM ammonium formate pH 3,75 in acetonitrile as solvent.

### Experimental infection of K18-hACE2 transgenic mice

Eight week-old K18-human ACE2 expressing C57BL/6 mice (B6.Cg-Tg(K18-hACE2)2Prlmn/J) were purchased from the Jackson Laboratory. For infection, mice (both sexes) were anesthetized by i.p. injection of ketamine (100 mg/kg) and xylazine (10 mg/kg) and then intranasally infected with 50 μl of DMEM containing (or not, in a mock sample) 5×10^2^ TCID_50_ of hCoV-19_IPL_France strain of SARS-CoV-2 (NCBI MW575140). Clofoctol (62.5 mg/kg in females and 50 mg/kg in males) was injected i.p. at 1h and 8h post-infection. The treatment was repeated the day after infection. Mice were sacrificed at day 2 or day 4 post-infection.

### Determination of viral loads in the lungs of mice

To determine the viral loads in lungs, half of right lobes were homogenized in Lysing Matrix D tubes (mpbio) containing 1 mL of PBS using Mixer Mill MM 400 (Retsch) (15min – 15 Hz). After centrifugation at 11,000 rpm for 5 min, the clarified supernatant was harvested for virus titration. Dilutions of the supernatant were done in DMEM with 1% penicillin/streptomycin and dilutions were transferred to Vero-E6 cells in 96-well plates for TCID_50_ assay. Quantitation of viral RNA in lung tissue was performed as follows. Briefly, half of the left lobe was homogenized in 1mL of RA1 buffer from the NucleoSpin RNA kit containing 20 mM of Tris(2-carboxyethyl)phosphine). Total RNAs in the tissue homogenate were extracted with NucleoSpin RNA from Macherey Nagel. RNA was eluted with 50μL of water.

### Determination of the viral load and assessment of gene expression by RT-qPCR

Half of the right lobe was homogenized in 1 mL of RA1 buffer from the NucleoSpin RNA kit containing 20 mM of TCEP. Total RNAs in the tissue homogenate were extracted with NucleoSpin RNA from Macherey Nagel. RNAs were eluted with 60 μL of water. RNA was reverse-transcribed with the High-Capacity cDNA Archive Kit (Life Technologies, USA). The resulting cDNA was amplified using SYBR Green-based real-time PCR and the QuantStudio™ 12K Flex Real-Time PCR Systems (Applied Biosystems™, USA) following manufacturers protocol. Relative quantifications were performed using the gene coding for RNA-dependent RNA polymerase (*RdRp*) and for glyceraldehyde 3-phosphate dehydrogenase (*Gapdh*). Specific primers were designed using Primer Express software (Applied Biosystems, Villebon-sur-Yvette, France) and ordered to Eurofins Scientifics (Ebersberg, Germany). The list of primers is available in Supplementary Table 3. Relative mRNA levels (2^−ΔΔCt^) were determined by comparing (a) the PCR cycle thresholds (Ct) for the gene of interest and the house keeping gene (ΔCt) and (b) ΔCt values for treated and control groups (ΔΔCt). Data were normalized against expression of the *gapdh* gene and are expressed as a fold-increase over the mean gene expression level in mock-treated mice. Viral load is expressed as viral RNA normalized to *Gapdh* expression level (ΔCt).

### Lung pathology scoring

Lung tissues were fixed in 4% PBS buffered formaldehyde for 7 days, rinsed in PBS, transferred in ethanol and then processed into paraffin-embedded tissues blocks. The subcontractor Sciempath Labo (Larçay, France) performed histological processing and analysis. The tissue sections in 3 μm were stained with haematoxylin and eosin (H&E) and whole mount tissues were scanned with a Nanozoomer (Hamatsu) and the morphological changes were assessed by a semi-quantitative score. For the scoring, a dual histopathology scoring system adapted from (Imai et al., 2020; Meyerholz and Beck, 2020) was used to assess pulmonary changes in mice. Inflammation was scored as 0 = absent, 1 = 1-10% of lung section, 2 = 11-25% of lung section, 3 = 26-50% of lung section, and 4=>50% of lung section affected.

### Statistical analysis

Results are expressed as the mean ± standard deviation (SD) unless otherwise stated. All statistical analysis was performed using GraphPad Prism v6 software. A Mann-Whitney *U* test was used to compare two groups unless otherwise stated. Comparisons of more than two groups with each other were analyzed with the One-way ANOVA Kruskal-Wallis test (nonparametric), followed by the Dunn’s posttest. *, P<0.05; **, P<0.01; ***, P<0.001.

### Ethics and biosafety statement

All experiments involving SARS-CoV-2 were performed within the biosafety level 3 facility of the Institut Pasteur de Lille, after validation of the protocols by the local committee for the evaluation of the biological risks and complied with current national and institutional regulations and ethical guidelines (Institut Pasteur de Lille/B59-350009). The experimental protocols using animals were approved by the institutional ethical committee “Comité d’Ethique en Experimentation Animale (CEEA) 75, Nord Pas-de-Calais”. The animal study was authorized by the “Education, Research and Innovation Ministry” under registration number APAFIS#25517-2020052608325772v3.

## Acknowledgements

We thank Sylvie van der Werf for sharing the SARS-CoV-2 strain BetaCoV/France/IDF0372/2020, Volker Thiel for providing HCoV-229E-RLuc and Chiesi for sharing clofoctol compound. We thank the infrastructure ChemBioFrance and the platforms ARIADNE-criblage (UMS2014-US41 PLBS) and ARIADNE-ADME to provide access to the Opera microscope and for LC-MS/MS analysis. Thank are also due to Nathan François for technical assistance and to Imène Belhaouane, Robin Prath and Nicolas Vandenabele for their technical help in the BSL3 facility. We are also grateful to Françoise Jacob-Dubuisson for her helpful comment on the manuscript. The immunofluorescence analyses were performed with the help of the imaging core facility of the BioImaging Center Lille Nord-de-France.

This work was supported by the Institut Pasteur de Lille, the Fondation pour la Recherche Médicale (FRM) and the Agence Nationale de la Recherche (ANR) (Project FRM_ANR Flash 20 ANTICOV), the Centre National de la Recherche Scientifique (CNRS: COVID and ViroCrib programs) and the I-Site Foundation (I-Site_Covid20_ANTI-SARS2). The platform used in this work was supported by the European Union (ERC-STG INTRACELLTB grant 260901), the ANR (ANR-10-EQPX-04-01), the “Fonds Européen de Développement Régional” (Feder) (12001407 [D-AL] EquipEx ImagInEx BioMed), CPER-CTRL (Centre Transdisciplinaire de Recherche sur la Longévité) and the Région Nord-Pas-de-Calais (convention 12000080).

## Author contributions

S.B., A.M., V.S., T.V., E.H., N.D., Y.R., Lo.D., A.D., C.R., K.S., L.B., C.M., C.P., A.B., A.V., Lu.D., Ju. D. and F.L. designed and/or performed experiments. S.B., A.M., V.S., T.V., E.H., N.D., Y.R., Lo.D., K.S., L.B., C.M., A.V., T.B. and S.H. analyzed data. I.E., E.K.A. and D.H. generated critical reagents. P.B., T.B., F.T., B.D. and Je.D. oversaw the conception and design of the experiments. S.B., T.V., T.B., F.T., and Je.D. wrote the manuscript.

---

**Supplementary Table S1.**
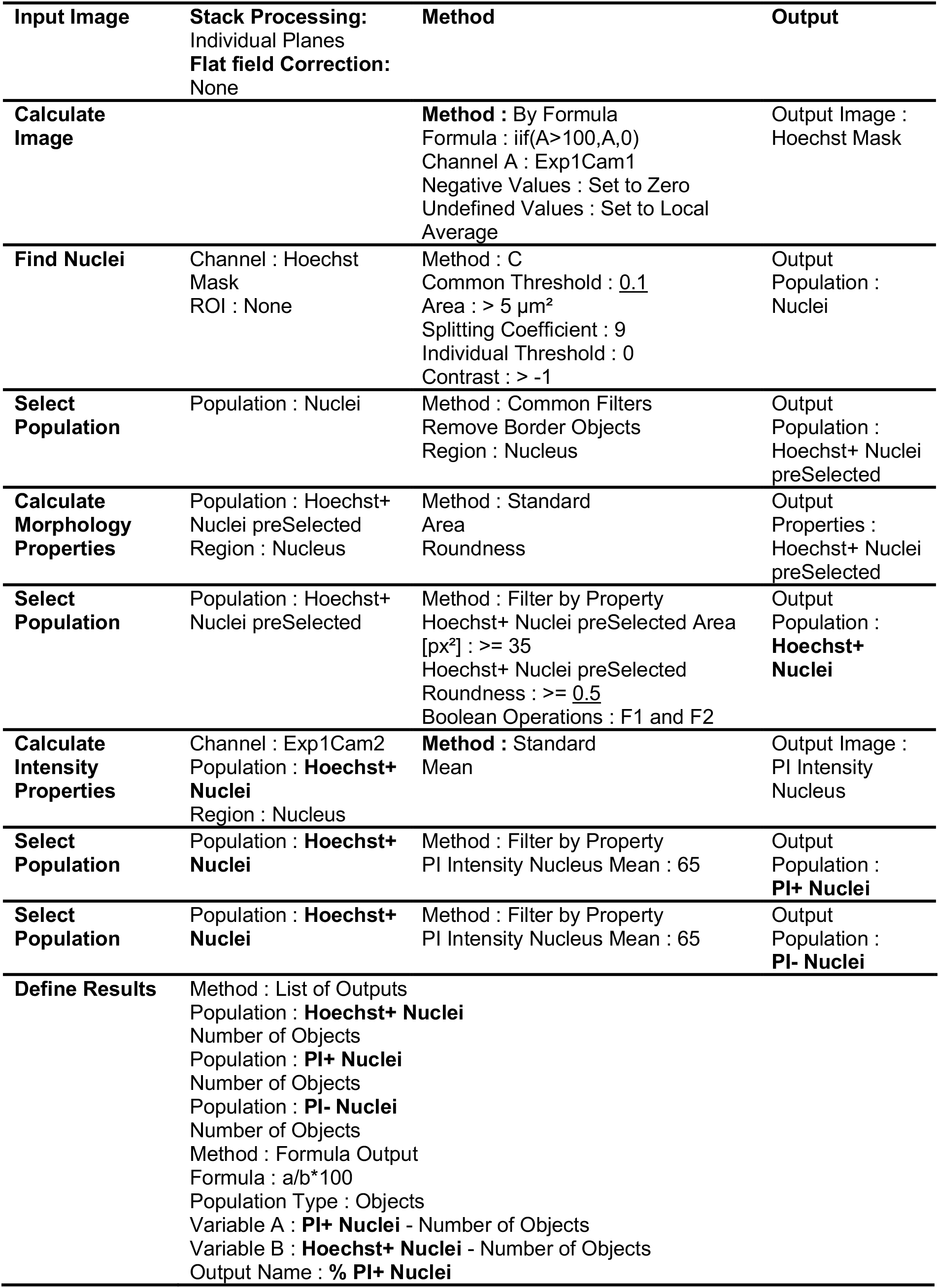
Multi-parametric script used in Columbus (version 2.3.1, PerkinElmer) for the determination of the percentage of PI+ Nuclei (PI-positive nuclei related to Fig. 1b).

| Input Image | Stack Processing:<br>Individual Planes<br>Flat field Correction:<br>None | Method | Output |
| --- | --- | --- | --- |
| Calculate Image | | Method : By Formula<br>Formula : $\text{iif}(A > 100, A, 0)$<br>Channel A : Exp1Cam1<br>Negative Values : Set to Zero<br>Undefined Values : Set to Local Average | Output Image :<br>Hoechst Mask |
| Find Nuclei | Channel : Hoechst Mask<br>ROI : None | Method : C<br>Common Threshold : <u>0.1</u><br>Area : $> 5 \mu\text{m}^2$<br>Splitting Coefficient : 9<br>Individual Threshold : 0<br>Contrast : $> -1$ | Output Population :<br>Nuclei |
| Select Population | Population : Nuclei | Method : Common Filters<br>Remove Border Objects<br>Region : Nucleus | Output Population :<br>Hoechst+ Nuclei preSelected |
| Calculate Morphology Properties | Population : Hoechst+ Nuclei preSelected<br>Region : Nucleus | Method : Standard<br>Area<br>Roundness | Output Properties :<br>Hoechst+ Nuclei preSelected |
| Select Population | Population : Hoechst+ Nuclei preSelected | Method : Filter by Property<br>Hoechst+ Nuclei preSelected Area [px<sup>2</sup>] : $\geq 35$<br>Hoechst+ Nuclei preSelected Roundness : $\geq \underline{0.5}$<br>Boolean Operations : F1 and F2 | Output Population :<br>**Hoechst+ Nuclei** |
| Calculate Intensity Properties | Channel : Exp1Cam2<br>Population : **Hoechst+ Nuclei**<br>Region : Nucleus | Method : Standard<br>Mean | Output Image :<br>PI Intensity Nucleus |
| Select Population | Population : **Hoechst+ Nuclei** | Method : Filter by Property<br>PI Intensity Nucleus Mean : 65 | Output Population :<br>**PI+ Nuclei** |
| Select Population | Population : **Hoechst+ Nuclei** | Method : Filter by Property<br>PI Intensity Nucleus Mean : 65 | Output Population :<br>**PI- Nuclei** |
| Define Results | Method : List of Outputs<br>Population : **Hoechst+ Nuclei**<br>Number of Objects<br>Population : **PI+ Nuclei**<br>Number of Objects<br>Population : **PI- Nuclei**<br>Number of Objects<br>Method : Formula Output<br>Formula : $a/b \times 100$<br>Population Type : Objects<br>Variable A : **PI+ Nuclei** - Number of Objects<br>Variable B : **Hoechst+ Nuclei** - Number of Objects<br>Output Name : % **PI+ Nuclei** | | |

**Figure S1:**
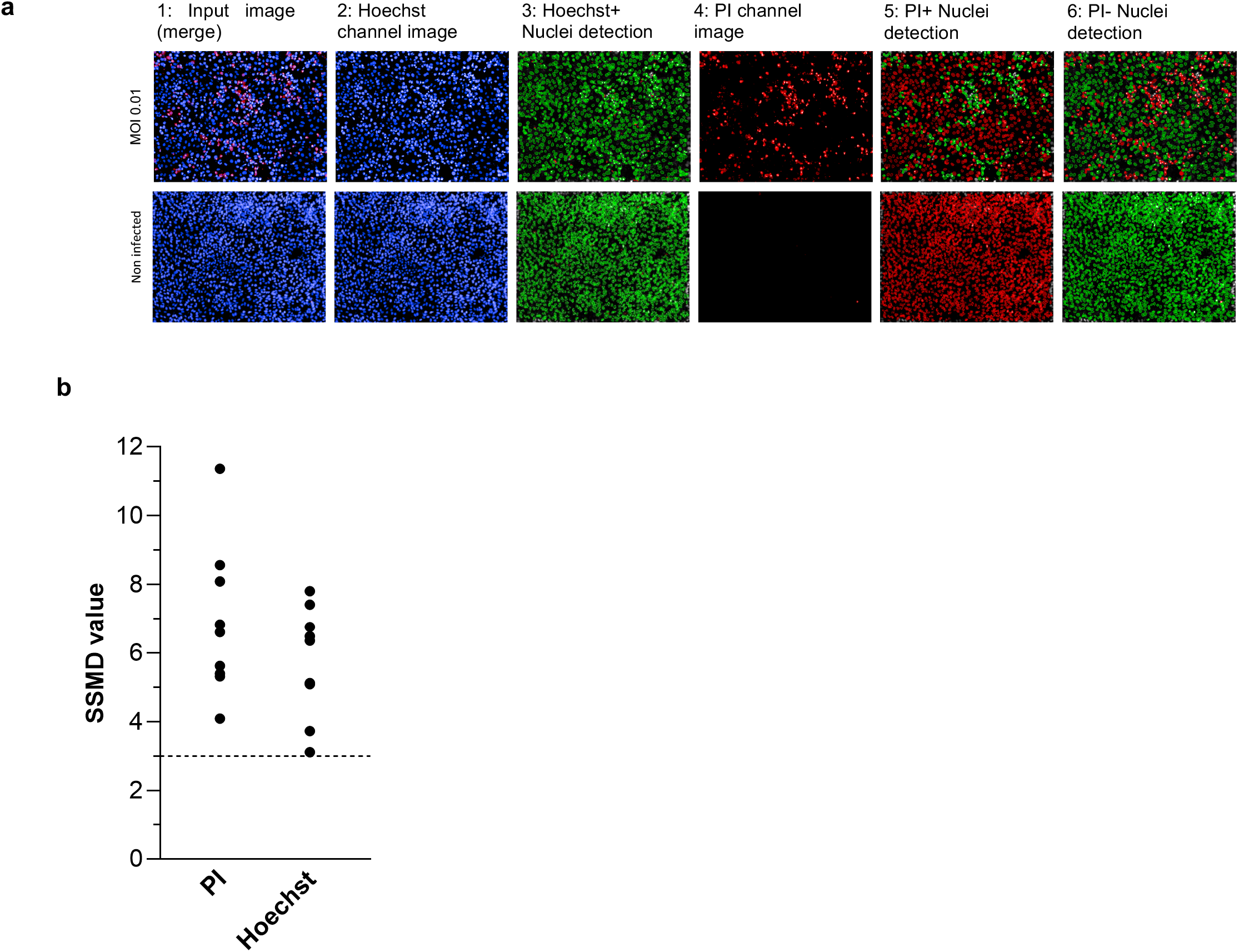
**a,** Typical images of Vero-81 cells infected with SARS-Cov-2 (top panel: MOI=0.01) or not (lower panel: Non-infected) acquired on an OPERA™ High Content Screening System (PerkinElmer) and corresponding image segmentation. 1: Typical 2-color images (Blue: Hoechst label, Red: PI-label); 2 and 4: 1-color images corresponding respectively to Hoechst and PI channel images; 3: Filled green objects correspond to total segmented nuclei, 5: Filled green objects correspond to PI positive cells or dead segmented cells, 6: Circled green cells correspond to non-infected cells. **b** SSMD values for HCS screen of the Apteeus TEELibrary® for the identification of anti-SARS-CoV-2 compounds. SSMD values were calculated for each of the 9 plates by comparing mean and standard deviations on both negative (Mock) and positive (Infected) controls. Dotted-line is indicative of a threshold of 3 allowing for the validation of the plates. SSMDs were calculated for both readouts.

**Figure S2:**
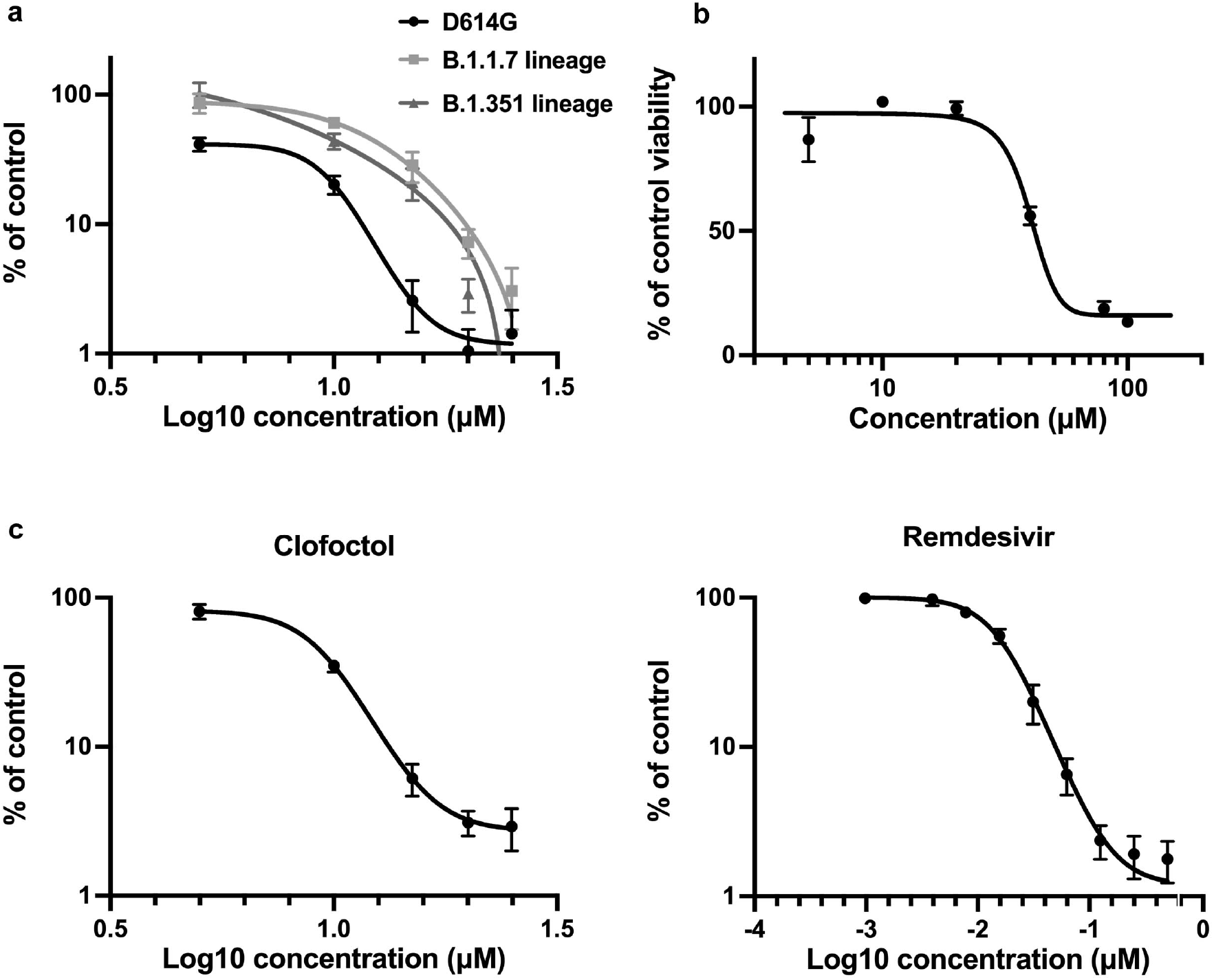
Clofoctol inhibits other SARS-CoV-2 variants as well as HCoV-229E. **a**, Vero-81 cells were infected either with SARS-CoV-2 of lineage B1 containing the D614G mutation (SARS-CoV-2/human/FRA/Lille_Vero-TMPRSS2/2020) or with SARS-CoV-2 of lineage B1.1.7 (GISAID accession number EPI_ISL_1653931) or lineage B.1.351 (GISAID accession number EPI_ISL_1653932). Viral genomes were quantified by RT-qPCR and normalized by the amount of total RNA. Results are presented as the percentage of the viral load of the control and represent the average of three independent experiments performed in duplicates. Error bars represent the standard error of the mean (SEM). **b**, Clofoctol is not cytotoxic in cell culture at concentrations below 40 μM. Huh-7 cells were cultured in the presence of given concentrations of clofoctol. Cell viability was monitored using the MTS-based viability assay after 24 hours of incubation. **c**, Huh-7 cells were infected with HCoV-229E-Rluc in presence of different concentrations of clofoctol or remdesivir. At 7h post-infection, cells were lysed and luciferase activities were quantified. Results are presented as the percentages of the control and represent an average of three independent experiments performed in triplicates. Errors bars represent the standard error of the mean (SEM).

**Figure S3:**
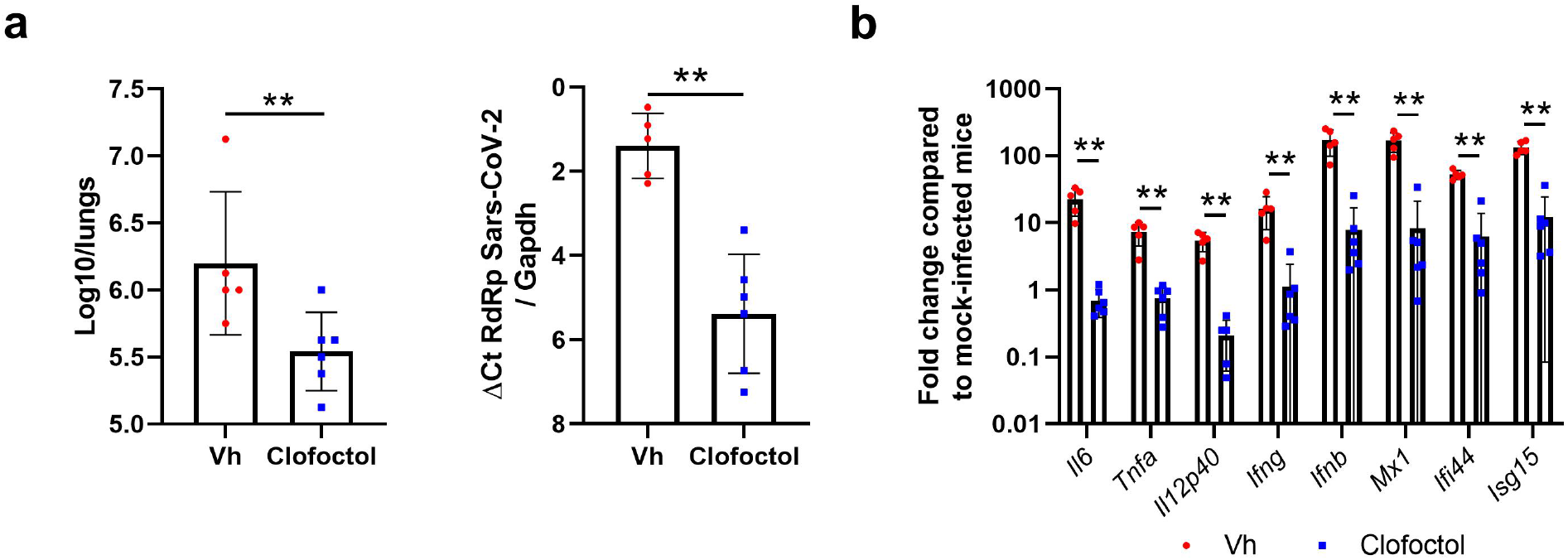
Effects of clofoctol treatment in male K18-hACE2 transgenic mice. **a** and **b**, Male mice were treated (50 mg/kg of clofoctol) and infected as described in Figure 4**b**. Mice were sacrificed at day 2 post-infection. **a**, The viral load was determined by titration on Vero-E6 cells (*left panel*) and by RT-qPCR (*right panel*). **b**, mRNA copy numbers of genes were quantified by RT-qPCR. Data are expressed as fold change over average gene expression in mock-treated (uninfected) animals. Results are expressed as the mean ± SD (n=5-6). Significant differences were determined using the Mann-Whitney U test (**p < 0.01).

**Figure S4:**
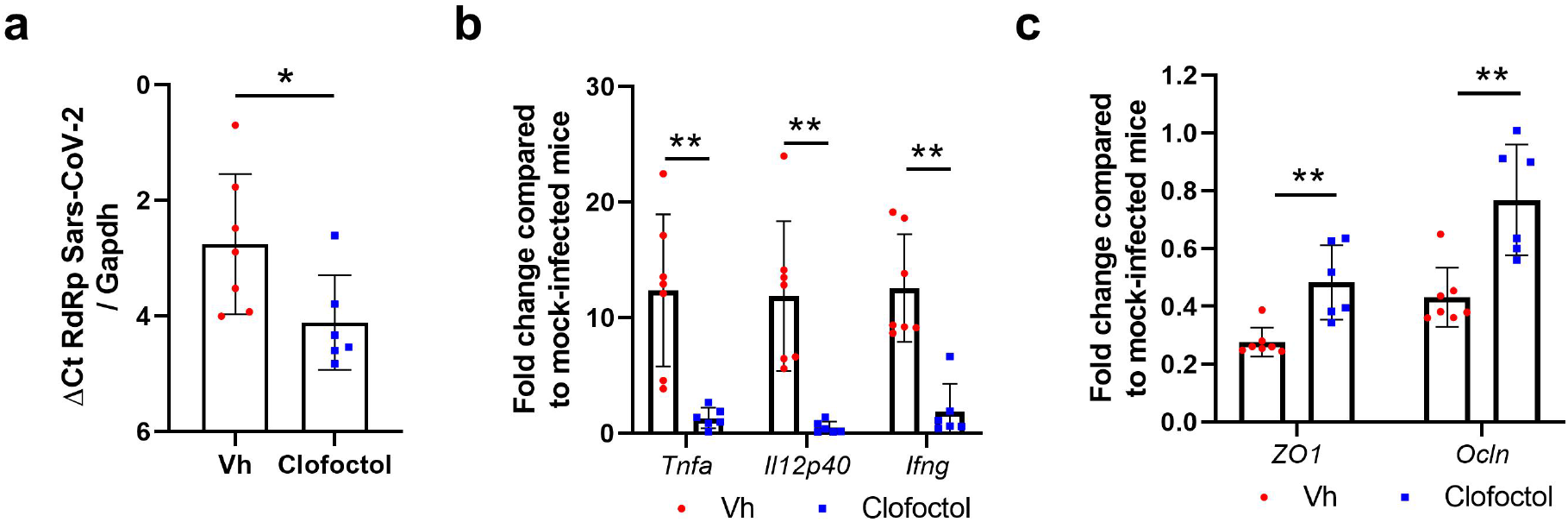
Effects of clofoctol treatment in female K18-hACE2 transgenic mice at day 4 post-infection. **a** and **b**, Female mice were treated and infected as described in Figure 4**b**. Mice were sacrificed at day 4 post-infection. mRNA copy numbers of genes were quantified by RT-qPCR. Panel **a** include inflammatory genes and panel **b** include genes involved in barrier function. Data are expressed as fold change over average gene expression in mock-treated (uninfected) animals. Results are expressed as the mean ± SD (n=6-7). Significant differences were determined using the Mann-Whitney U test (*p < 0.05; **p < 0.01).

